# Cerebellar Morphological Differences in Bipolar Disorder Type I

**DOI:** 10.1101/2023.02.20.528549

**Authors:** Gail I. S. Harmata, Ercole John Barsotti, Lucas G. Casten, Jess G. Fiedorowicz, Aislinn Williams, Joseph J. Shaffer, Jenny Gringer Richards, Leela Sathyaputri, Samantha L. Schmitz, Gary E. Christensen, Jeffrey D. Long, Marie E. Gaine, Jia Xu, Jake J. Michaelson, John A. Wemmie, Vincent A. Magnotta

**Affiliations:** Department of Psychiatry, The University of Iowa, United States; Iowa Neuroscience Institute, The University of Iowa, United States; Department of Epidemiology, The University of Iowa, United States; Interdisciplinary Graduate Program in Genetics, The University of Iowa, United States; Department of Electrical and Computer Engineering, The University of Iowa, United States; Department of Radiation Oncology, The University of Iowa, United States; Department of Radiology, The University of Iowa, United States; Department of Biomedical Engineering, The University of Iowa, United States; Department of Biostatistics, University of Iowa, The University of Iowa, United States; Department of Pharmaceutical Sciences and Experimental Therapeutics (PSET), College of Pharmacy, The University of Iowa, United States; Department of Molecular Physiology and Biophysics, The University of Iowa, United States; Department of Neurosurgery, The University of Iowa, United States; Department of Biosciences, Kansas City University, United States; Department of Psychiatry, University of Ottawa, Canada; Veterans Affairs Medical Center, Iowa City, United States

**Keywords:** Bipolar Disorder, Cerebellum, Cerebellar Volume, Mood Disorders

## Abstract

**Background:** The neural underpinnings of bipolar disorder (BD) remain poorly understood. The cerebellum is ideally positioned to modulate emotional regulation circuitry yet has been understudied in BD. Previous studies have suggested differences in cerebellar activity and metabolism in BD, however findings on cerebellar structural differences remain contradictory.

**Methods:** We collected 3T anatomical MRI scans from participants with (N = 131) and without (N = 81) BD type I. Differences in cerebellar volumes were assessed along with factors that influence the results.

**Results:** The cerebellar cortex was smaller bilaterally in participants with BD. Polygenic propensity score (bipolar N = 103, control N = 64) did not predict any cerebellar volumes, suggesting that non-genetic factors may have greater influence on the cerebellar volume difference we observed in BD. Cerebellar white matter volumes increased with more adverse childhood events, but we did not observe any associations with parental psychiatric illness. We also evaluated time from onset and symptom burden and found no associations with cerebellar volumes, suggesting neurodevelopment may differ prior to onset. Finally, we found taking sedatives was associated with larger cerebellar white matter and non-significantly larger cortical volume.

**Limitations:** This study was cross-sectional, limiting interpretation of possible mechanisms. Most of our participants were White, which could limit the generalizability. Additionally, we did not account for potential polypharmacy interactions.

**Conclusions:** These findings suggest that external influences, such as medications, may influence cerebellum structure in BD and may mask underlying differences. Accounting for medication may be critical for consistent findings in future studies.

## Introduction

Bipolar disorder type I is a chronic condition characterized by mood instability that affects an estimated 1% of the U.S. population (Clemente et al., 2015). Bipolar disorder is often debilitating, disrupting personal, work-related, and social function (Alonso et al., 2011). Despite its seriousness, the cause of bipolar disorder remains unknown. A number of brain imaging studies have been conducted to gain insight into the underlying neurobiology by examining brain morphology, function, and metabolism. Although there is variability in the literature related to morphological changes associated with bipolar disorder, several meta- and mega-analyses have observed reduced overall cerebral volume and increased ventricle size in bipolar disorder along with more specific volume changes of the prefrontal cortex, cingulate cortex, temporal lobe, and subcortical structures such as the striatum and amygdala (de Zwarte et al., 2019; Hallahan et al., 2011; Houenou et al., 2011; Wise et al., 2017). Functional differences have similarly been observed in overlapping brain regions (Chen et al., 2022). These identified areas are consistent with a model of dysregulated emotional control in bipolar disorder proposed by Strakowski and colleagues (Strakowski et al., 2012). However, the exact nature of such dysregulation is not precisely understood.

One brain area of potential importance to understanding emotional dysregulation in bipolar disorder is the cerebellum. Although often associated with coordination and motor control, the cerebellum has been increasingly recognized to contribute to cognitive functioning and possibly psychiatric disorders (Phillips et al., 2015; Schmahmann, 2019). The cerebellum has connections to nodes of the emotional control network, including the amygdala and prefrontal cortex (Blatt et al., 2013). Several reports have implicated the cerebellum in bipolar disorders. Studies have observed altered metabolism and neurochemistry (Cecil et al., 2003; Lai et al., 2019; Magnotta et al., 2022; Wu et al., 2021); differences in T1ρ relaxation times, potentially reflecting pH differences (Johnson et al., 2018; Johnson et al., 2015); and altered functional connectivity (Cui et al., 2022; Shinn et al., 2017). Furthermore, cerebellar activity has been linked to mood state in participants with bipolar disorder (Fleck et al., 2011; Shaffer et al., 2018), supporting a potential model of cerebellar involvement in mood regulation.

Findings from structural imaging studies of the cerebellum in bipolar disorder have reported inconsistent findings. While many studies have found the cerebellum or regions thereof (e.g., hemispheric gray matter, vermal subregions, etc.) to be smaller in bipolar disorder compared to controls (Baldaçara et al., 2011; Berk et al., 2017; de Zwarte et al., 2019; Kim et al., 2013; Lin et al., 2018; Lisy et al., 2011; Moorhead et al., 2007; Rimol et al., 2010; Sarıçiçek et al., 2015), other studies have found the opposite (Adler et al., 2007; Cui et al., 2011; Wise et al., 2017) or no differences (Demirgören et al., 2019; Goikolea et al., 2018; Laidi et al., 2015; McDonald et al., 2005). One possible explanation for some of these contradictory results may be explained by differences in sampling time points: some have found smaller or non-significantly smaller cerebellar volumes in bipolar disorder associated with age or number of mood episodes (DelBello et al., 1999; Lisy et al., 2011; Monkul et al., 2008; Moorhead et al., 2007; Yates, 1987). This could indicate that the cerebellum experiences accelerated atrophy with the disorder. Furthermore, one study found greater reduction in cerebellar vermis gray matter density when accounting for medication status (Kim et al., 2013), while another study reported larger cerebellar cortex and white matter volumes with lithium treatment (Jones et al., 2022); thus, medication effects make interpretation of findings in bipolar disorder challenging. In addition, genetic loading for bipolar disorder may also influence cerebellar volumes. For example, a prior study by Brambilla et al. found that participants with bipolar disorder and a family history of mood disorders had smaller volumes than participants with bipolar disorder without a family history (Brambilla et al., 2001). Another study found that both participants with bipolar disorder and their first-degree relatives had smaller cerebellar volumes than controls (Sarıçiçek et al., 2015; but see de Zwarte et al., 2019). Finally, many bipolar disorder studies have had relatively small sample sizes or did not distinguish between bipolar disorder subtypes. Some or all of these factors may have contributed to the heterogeneity of the results that have been reported in the literature.

The lack of consensus in the literature makes understanding the nature of cerebellar involvement in bipolar disorder challenging. To address this issue, we examined cerebellar morphology in a large single-site sample composed of participants with bipolar disorder type I (n = 131) and a comparison control sample (n = 81). We hypothesized that we would find significantly smaller cerebellar volumes in participants with bipolar disorder compared to controls, consistent with the largest multi-site study (de Zwarte et al., 2019). We also hypothesized that genetic loading for bipolar disorder would be associated with the volume of the cerebellum based on the reported differences in affected participants with family history of bipolar disorder (Brambilla et al., 2001) and in unaffected relatives (de Zwarte et al., 2019; Sarıçiçek et al., 2015). Finally, we evaluated whether potentially contributing variables such as adverse childhood events, time from mood symptom onset, self-reported symptom burden, and medication status would correlate with cerebellar volumes.

## Methods

### Participants

Following University of Iowa Institutional Review Board approval, individuals with bipolar disorder type I and control participants were recruited to participate into a neuroimaging and genetics study (Fig. 1) after providing written informed consent. All possible participants were screened for exclusion criteria, including neurological comorbidities, current drug or alcohol abuse, prior loss of consciousness for more than 10 minutes, or MRI contraindications. A mid-study exclusion criterion for controls currently taking a medication for depression, schizophrenia, or bipolar disorder was applied retroactively, which excluded one previously collected control participant from the analyses who was receiving active treatment for depression. Controls were recruited in an attempt to frequency-match to participants with bipolar disorder type I (subsequently referred to as simply bipolar disorder in the remainder of the text) based on age, sex, race, and subjective social economic status from self-report on the MacArthur Scale of Subjective Social Status, on which participants identify their relative standing on a 10-rung ladder (Adler et al., 2000). After obtaining informed written consent, participants were interviewed and participants with bipolar disorder had their diagnosis confirmed via the Structured Clinical Interview for DSM Disorders (SCID). Additional information was also collected, including the Montgomery-Åsberg Depression Rating Scale (MADRS), Young Mania Rating Scale (YMRS), the Beck Anxiety Inventory (BAI), Adverse Childhood Event (ACE) Score, parental psychiatric history, and a list of current medications. Psychiatric medications were sorted into classes modified from the WHO Anatomical Therapeutic Chemical system (WHO Collaborating Centre for Drug Statistics Methodology, 2022); the drug classes included antidepressants, antipsychotics, sedatives, anticonvulsants, stimulants, and lithium. Participants with bipolar disorder were asked to retrospectively report estimated symptom burden over the course of the last decade (or since diagnosis, if less than 10 years prior), an approach used previously (Sodhi et al., 2012). Finally, participants had a blood sample taken for the purpose of genetic analysis, with 188 samples processed at the time of evaluation.

**Figure 1.**
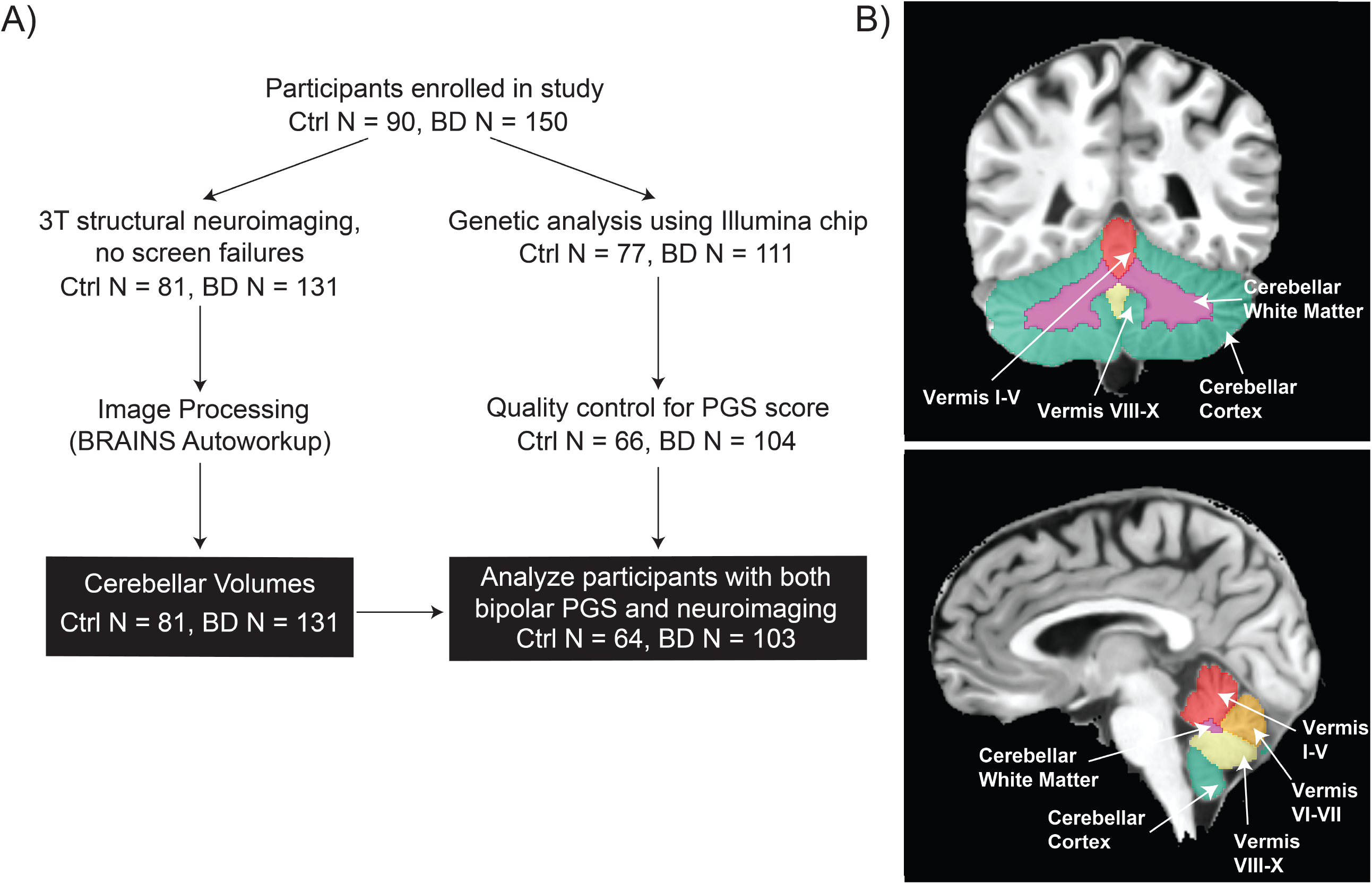
Study Schematic. (A) We recruited 150 participants with bipolar disorder type I and 90 control participants. After excluding screen failures and participants that were unable to complete the 3T anatomical scan, we obtained cerebellar volumetric data from most participants (81 control participants, 130 bipolar participants). During the site visit, we also collected blood samples. As of the time of this study, 64 control participants and 103 bipolar participants had both anatomical scans and polygenic risk scores calculated from the blood sample analysis. (B) Coronal and sagittal views of cerebellar segmentation used for most analyses in this report. This segmentation resulted from a custom atlas based on the Neuromorphometrics segmentation (Neuromorphometrics Inc., 2007).

**Figure 2.**
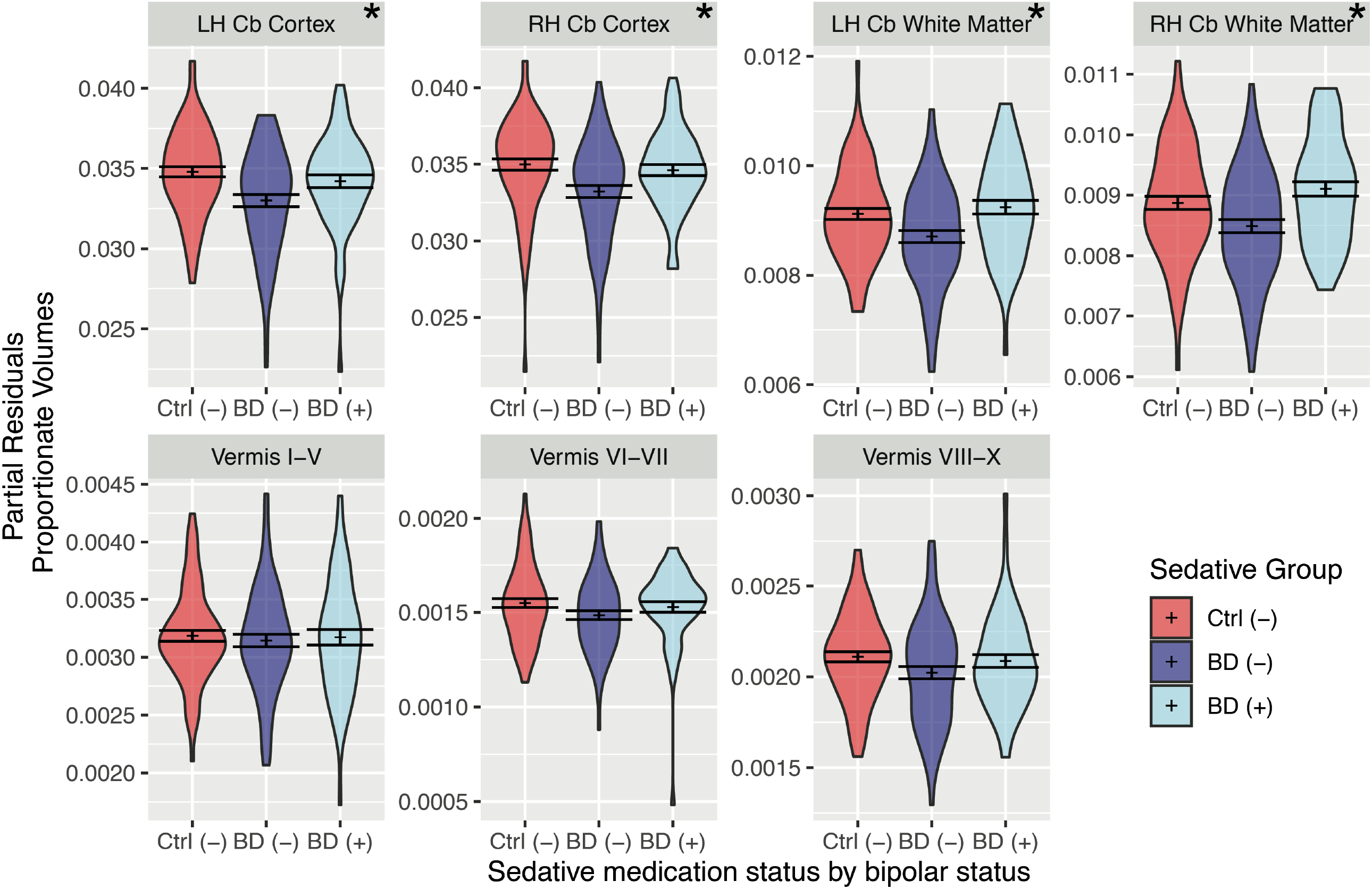
Bipolar participants taking sedative medication (BD (+), N = 56) had similar cerebellar white matter and gray matter volumes to control participants not on sedative medication (Ctrl (-), N = 75), while bipolar participants not taking sedatives generally had smaller volumes than the other groups (BD (-), N = 75). Graphs show partial residuals for diagnosis/medication group variable from models controlling for age and sex. Regions with significant effect of diagnosis/medication group variable (q < 0.05) are marked with asterisks (*).

### 3T MR Imaging Protocol

Imaging was conducted on either a GE 750W or Premier 3T scanner with a 32 channel or 48 channel head coil, respectively. The same imaging protocol was acquired across both scanner configurations, which included the following anatomical imaging scans: 3D coronal T1-weighted BRAVO (TE=2.3ms, TR=6.8ms, TI=450ms, Flip Angle=12°, FOV=256×256×220mm, Matrix=256×256×220, Bandwidth=488 Hz/Pixel, Parallel Imaging Factors =2 phase x 1 slice) and 3D sagittal T2-weighted CUBE (TE=98ms, TR=3202ms, FOV=256×256×200, Matrix=256×256×200, Bandwidth=488 Hz/Pixel, Parallel Imaging Factors =2 phase x 2 slice). Additional MR scans were also collected including 3D T1ρ imaging, resting state fMRI, and multi-shell diffusion imaging data, but were not used in this study. Most participants also completed a 7T metabolic imaging session as previously reported in Magnotta et al. (2022).

### MRI Image Analysis

The T1 and T2 weighted scans described above were analyzed using BRAINS AutoWorkup (Forbes et al., 2016; Ghayoor et al., 2018; Kim et al., 2015; Kim et al., 2016), which includes the following steps: 1) AC-PC alignment; 2) bias field correction; 3) tissue classification; and 4) anatomical labeling using a custom atlas based upon the Neuromorpometrics segmentation (Neuromorphometrics Inc., 2007). This atlas includes parcellation of the cerebral cortex, white matter, subcortical structures, brainstem, and cerebellum. The cerebellum was divided into gray matter and white matter as well as vermis lobules (I-V, VI-VII, and VIII-X) (**Figure 1B**). All subdural tissue components were combined to generate an estimate of intracranial volume. Regional volumetric data was divided by intracranial volume (ICV) to control for differences in cranial size. This resulted in proportional volumetric data. For these analyses, we focused on the cerebellar regions of interest which included left/right cerebellar gray matter, left/right cerebellar white matter, and three cerebellar vermis lobules (I-V, VI-VII, and VIII-X). To explore cerebellar volume differences in more detail, we also parcellated the cerebellum using the SUIT atlas (Diedrichsen et al., 2009) by non-linear registration of the MNI atlas to the AC-PC aligned image for each participant using ANTS (Avants et al., 2008). The resulting transformation was applied to the SUIT atlas resampled using nearest neighbor interpolation, which divided the cerebellum into the following cerebellar subdivisions: I-IV, V, VI, Crus I, Crus II, VIIb, VIIIa, VIIIb, IX, and X. These levels were divided into left and right hemispheres, and vermal subdivisions were available for VI-X.

### SNP genotyping and quality control

Available samples were genotyped with Illumina’s Infinium PsychArray-24 v1.3 BeadChip (n = 188), genotypes were called using GenomeStudio2.0 software with Illumina’s provided cluster files. PLINK (Purcell et al., 2007) was used for quality control. Data was filtered down to SNPs, removing indels and non-biallelic sites. SNPs with low minor allele frequency in our sample (< 5%) were removed. All samples had high genotyping rates (> 99%). SNPs defying Hardy-Weinberg equilibrium were removed (p-value < 1 × 10^−6^). Four individuals were removed due to relatedness (identity-by-descent cutoff of 0.125), 8 were removed due to high heterozygosity rates, and 6 more were removed due to population structure (i.e., did not cluster with the European superpopulation in 1000 Genomes (1000 Genomes Project Consortium et al., 2015)).

The final sample size for the genetic analysis was 170, with data available from 104 participants with bipolar disorder and 66 controls. Data for the 170 individuals with 350,345 SNPs passing quality control were then imputed to the HRC r1.1 hg19 build reference dataset (Loh et al., 2016) using Michigan Imputation Server MiniMac4 (Das et al., 2016). SNPs with imputation *R*^2^ scores < 0.3 were dropped. Imputed genotype dosages were converted to hard calls using a cutoff of 90% probability. Finally, SNPs were subset to HapMap3 SNPs (International HapMap 3 Consortium et al., 2010) for subsequent polygenic propensity score calculations (final number of SNPs = 1,053,095).

### Polygenic Propensity Scores (PGS)

GWAS summary statistics from the most recent PGC bipolar disorder study (Mullins et al., 2021) were used to compute PGS in the 170 samples with the quality-controlled genotype and neuroimaging data. PGS were calculated for our cohort using LDpred2 (Privé et al., 2020) implemented by the “bigsnpr” package (Privé et al., 2018) in R (R Core Team, 2021). For linkage disequilibrium reference, a dataset of 362,320 European individuals of the UK Biobank (provided by the developers of LDpred2) was used for calculating the genetic correlation matrix, estimating heritability, and calculating infinitesimal beta weights. PGS were then corrected for population stratification and z-scaled by sex. Of the 170 participants with a calculated PGS, 166 participants (103 cases and 64 controls) had available neuroimaging data and met all other inclusion criteria for this report as described above.

### Statistical Analysis

Analysis was performed in R version 4.1.1 (R Core Team, 2021) and RStudio (RStudio Team, 2020) using the following packages: car (Fox and Weisberg, 2019), arsenal (Heinzen et al., 2021), emmeans (Lenth, 2022), and tidyverse (Wickham et al., 2019). We tested cerebellar white matter, cerebellar gray matter, and vermal volumes separately using linear regression models to assess whether diagnosis with bipolar disorder significantly predicted volume. Next, we tested the effects of genetic loading for bipolar disorder on cerebellar volumes while employing the same covariates by replacing diagnosis in the linear models with PGS for bipolar disorder. We followed these analyses by evaluating the data only from the participants with bipolar disorder to determine if parental history, adverse childhood events, duration of illness, symptom burden (percent time manic, depressed, or euthymic), or medication status predicted cerebellar volumes. For the medication follow-up analysis, we used ANCOVA models with Tukey-corrected pairwise comparisons to assess the effect of the following groupings: control participants not on the medication class, Ctrl (-); participants with bipolar disorder not on the medication class, BD (-); and participants with bipolar disorder on the medication class, BD (+). To adjust for the effects of age in years (linear effect) and sex (categorical effect), we included these as covariates in all linear regression and ANCOVA models. To correct for testing multiple cerebellar regions, we utilized false discovery rate (FDR) correction; the resulting q-values were considered significant if q < 0.05. Demographic differences (**Table 1**) were assessed by t-test or chi-square test as appropriate.

**Table 1.**
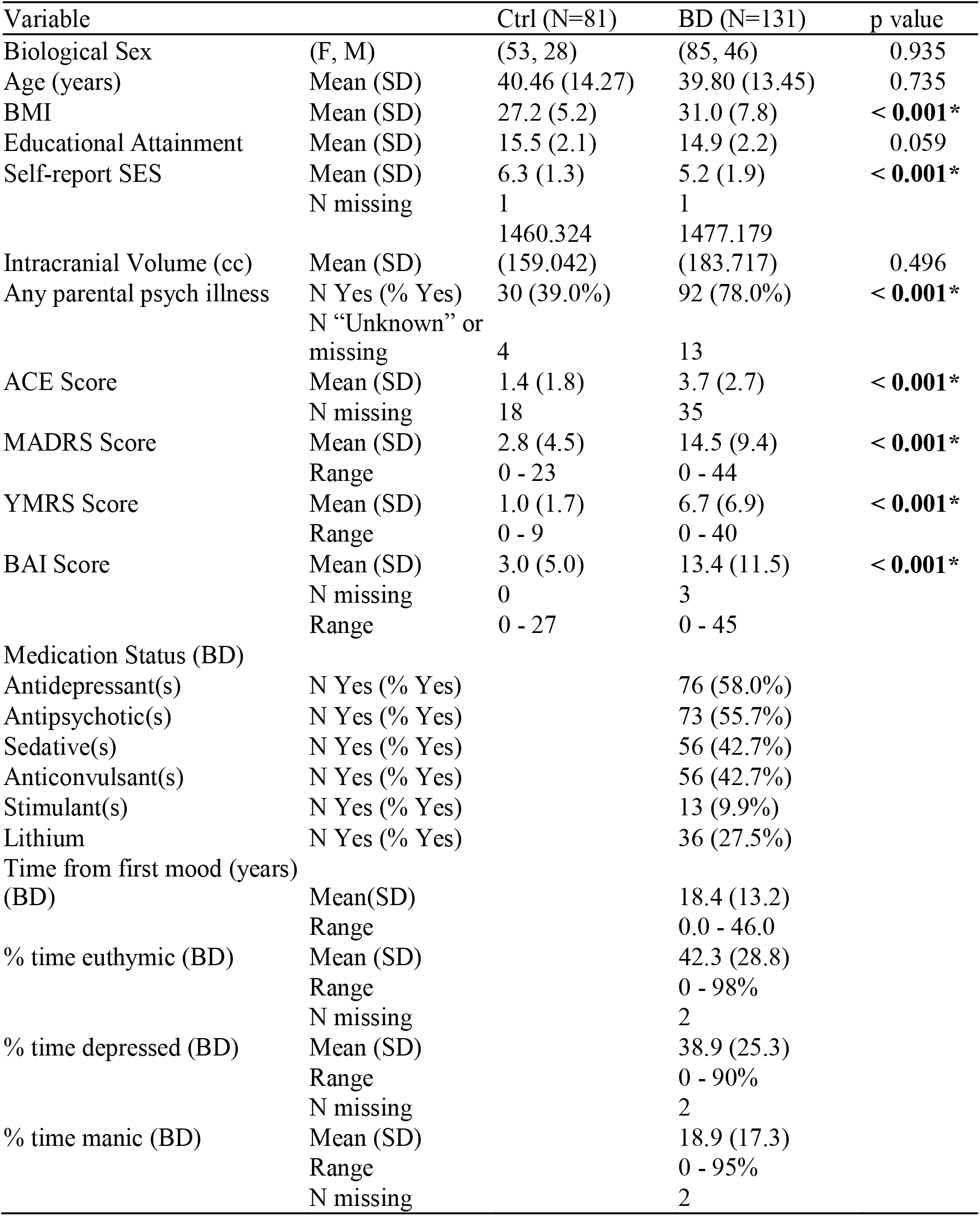
Demographics for control participants (Ctrl, N = 81) and bipolar type I (N = 131) participants (BD). Significant differences are indicated (p < 0.05, *).

**Table 2.** Cerebellar cortex volumes are smaller bilaterally in bipolar disorder type I. A total of 211 participants (81 Ctrl, 131 BD) were included in the analysis, and age and sex were used as covariates. P-values were FDR-corrected. Significant differences are indicated (q < 0.05, *).

| Cerebellar Volume<br>(proportion of<br>ICV) | Effect of bipolar diagnosis<br>(N “Ctrl” = 81, N “BD” = 131) |  |  |  |
| --- | --- | --- | --- | --- |
| | Beta Coefficient<br>Estimate (SE) | Estimated<br>Group Difference<br>(converted to $\text{cm}^3$) | p-value | q-value |
| Left Cortex | -1.388e-03 (4.333e-04) | -2.042 | 0.002 | **0.011\*** |
| Right Cortex | -1.273e-03 (4.540e-04) | -1.872 | 0.006 | **0.019\*** |
| Left White Matter | -2.244e-04 (1.326e-04) | -0.330 | 0.092 | 0.129 |
| Right White Matter | -1.501e-04 (1.349e-04) | -0.221 | 0.267 | 0.312 |
| Vermis I-V | -3.757e-05 (6.434e-05) | -0.055 | 0.560 | 0.560 |
| Vermis VI-VII | -5.276e-05 (2.929e-05) | -0.078 | 0.073 | 0.129 |
| Vermis VIII-X | -6.558e-05 (3.823e-05) | -0.096 | 0.088 | 0.129 |

## Results

### Demographics

The demographics for our sample are shown in **Table 1**. The control and bipolar disorder groups were not significantly different in age, sex, and intracranial volume (all p > 0.05). However, participants with bipolar disorder had a non-significantly lower educational attainment (p = 0.059), and participants with bipolar disorder had significantly lower perceived socioeconomic status than controls (5.2 vs 6.3, p < 0.001) and a higher BMI (31.0 v. 27.2 kg/m^2^, p < 0.001). In addition, participants with bipolar disorder more frequently reported a parental history of psychiatric illness (78.0% vs. 39.0%, p < 0.001) and a greater number of adverse childhood events (3.7 vs. 1.4 ACEs; p < 0.001) than controls. It is important to note that household mental illness is considered an ACE and may contribute to other factors included in the overall ACE score such as parental separation or drug/alcohol abuse in the household.

Finally, participants with bipolar disorder had significantly higher depression, mania, and anxiety ratings (all p < 0.001).

Within the participants with bipolar disorder, the sample included a wide range of time from first mood episode (0 to 46 years), with an average of 18.4 years since onset. Similarly, we were able to capture a variety of self-reported symptom burdens, with ranges of 0% to more than 90% of time spent in each mood state over the past decade (or since diagnosis if less than 10 years ago). On average, participants with bipolar disorder spent a similar amount of time euthymic and depressed states (42.3% and 38.9% respectively), with a lower amount of time in a manic state (18.9%).

### Differences in cerebellar volume in bipolar disorder

To investigate differences in cerebellar volume in bipolar disorder, we conducted MRI imaging and analyzed the resulting cerebellar data, normalized to intracranial volume, using linear regression controlling for age and sex (**Table 2**). We found that participants with bipolar disorder had smaller cerebellar gray matter volumes compared to controls with estimated mean differences of -2.042 cm^3^ (p = 0.002, q = 0.011) and -1.872 cm^3^ (p = 0.006, q = 0.019) for left and right hemispheres, respectively. We did not detect significant differences in cerebellar white matter volumes nor vermal volumes (all p > 0.05 and all q > 0.05), but all cerebellar regions tended to be smaller in participants with bipolar disorder. To determine whether the observed differences in cerebellar gray matter were driven by a specific region, we used the SUIT atlas (Diedrichsen et al., 2009) to parcellate the cerebellum and explored all resulting volumes. Numerous hemispheric regions were significantly or non-significantly smaller in bipolar disorder (**Supplemental Table 1, Supplemental Fig. 1**), suggesting differences in cerebellar volume are diffuse throughout much of the cerebellum.

### Association between genetic loading for bipolar disorder cerebellar volume

We subsequently explored whether genetic loading for bipolar disorder would be associated with cerebellar volume. First, we calculated polygenic propensity score (PGS) for bipolar disorder for all participants with available data (N = 103 bipolar disorder and N = 64 control). As expected, the participants with a diagnosis of bipolar disorder had significantly higher bipolar PGS than the control group (t-test, t(165) = -4.856, p < 0.001; **Supplemental Figure 2**), suggesting that this summary measure of genetic loading does have predictive value for bipolar disorder. Next, we tested whether bipolar PGS would predict any cerebellar volumes across both diagnosis groups, adjusting for age and sex (**Table 3**). Bipolar PGS did not predict any cerebellar volumes (all p > 0.05 and all q > 0.05). This suggests that overall genetic loading may not explain the observed differences in gray matter volumes between participants with bipolar disorder and controls.

**Table 3.** Bipolar polygenic propensity score (PGS) was not associated with differences in cerebellar volumes. A total of 167 participants (64 Ctrl, 103 BD) had available bipolar PGS and structural scans and were included in the analysis. Age and sex were used as covariates. P-values were FDR-corrected.

| Cerebellar Volume<br>(proportion of ICV) | Effect of bipolar PGS (N = 167) |  |  |
| --- | --- | --- | --- |
|  | Beta Coefficient<br>Estimate (SE) | p-value | q-value |
| Left Cortex | -4.000e-04 (2.367e-04) | 0.093 | 0.213 |
| Right Cortex | -3.793e-04 (2.345e-04) | 0.108 | 0.213 |
| Left White Matter | -9.530e-05 (6.916e-05) | 0.170 | 0.238 |
| Right White Matter | -1.063e-04 (6.835e-05) | 0.122 | 0.213 |
| Vermis I-V | -8.395e-06 (3.433e-05) | 0.807 | 0.807 |
| Vermis VI-VII | 1.771e-05 (1.644e-05) | 0.283 | 0.330 |
| Vermis VIII-X | -3.277e-05 (1.991e-05) | 0.102 | 0.213 |

### Effect of adverse childhood events on cerebellar volumes in bipolar disorder

To better understand what may contribute to the variation in cerebellar volumes within participants with bipolar disorder, we explored whether either parental history of psychiatric disorders or number of adverse childhood events (ACE score) was associated with cerebellar volumes (**Table 4**). As previously mentioned, ACE score is not independent from familial history of psychiatric illness. In addition, familial history of psychiatric illness also contains a genetic contribution. With these limitations in mind, parental psychiatric history did not predict any cerebellar volumes (N = 118; all p > 0.05 and q > 0.05), while ACE score did significantly predict volumes of left cerebellar gray matter (N = 96; p = 0.019, q = 0.033), right cerebellar gray matter (p = 0.018, q = 0.033), left cerebellar white matter (p = 0.003, q = 0.019), and right cerebellar white matter (p = 0.009, q = 0.031). Thus, it is possible that cerebellar growth may be responsive to environmental influences. However, further work is needed to tell whether this represents a true distinction between genetics versus formative experiences. With higher ACE scores, all cerebellar white and gray matter volumes tended to be higher, which was contrary to what was expected.

**Table 4.** Within participants with bipolar disorder, parental history of psychiatric illness was not associated with differences in cerebellar volumes; however, adverse childhood experience (ACE) score was associated with larger cerebellar cortex and white matter volumes bilaterally. These variables were tested in separate models with age and sex as covariates. P-values were FDR-corrected. Significant results are indicated (q < 0.05, *).

| Variable of interest<br>(BD only) | Cerebellar Volume<br>(prop. of ICV) | Beta<br>Coefficient<br>Estimate (SE) | p-value | q-value |
| --- | --- | --- | --- | --- |
| Parental Psychiatric<br>History<br>(N “Yes” = 92,<br>N “No” = 26) | Left Cortex | 3.373e-04<br>(7.207e-04) | 0.641 | 0.787 |
|  | Right Cortex | 3.002e-04<br>(7.136e-04) | 0.675 | 0.787 |
|  | Left White Matter | 2.038e-04<br>(2.228e-04) | 0.362 | 0.787 |
|  | Right White Matter | 1.232e-04<br>(2.190e-04) | 0.575 | 0.787 |
|  | Vermis I-V | 1.372e-04<br>(1.092e-04) | 0.211 | 0.787 |
|  | Vermis VI-VII | -2.174e-05<br>(4.811e-05) | 0.652 | 0.787 |
|  | Vermis VIII-X | -5.834e-06<br>(6.355e-05) | 0.927 | 0.927 |
| ACE Score<br>(N = 96) | Left Cortex | 2.839e-04<br>(1.189e-04) | 0.019 | **0.033\*** |
|  | Right Cortex | 2.807e-04<br>(1.170e-04) | 0.018 | **0.033\*** |
|  | Left White Matter | 1.144e-04<br>(3.709e-05) | 0.003 | **0.019\*** |
|  | Right White Matter | 1.001e-04<br>(3.735e-05) | 0.009 | **0.031\*** |
|  | Vermis I-V | 3.693e-05<br>(1.796e-05) | 0.043 | 0.060 |
|  | Vermis VI-VII | 1.442e-05<br>(8.065e-06) | 0.077 | 0.090 |
|  | Vermis VIII-X | 1.012e-05<br>(1.117e-05) | 0.367 | 0.367 |

### Association between duration of illness and symptom burden with cerebellar volumes

Since previous work has suggested that an increased number of episodes or illness duration may be associated with changes in cerebellar volumes in bipolar disorder (DelBello et al., 1999; Lisy et al., 2011; Monkul et al., 2008; Moorhead et al., 2007; Yates, 1987), we tested whether duration of illness or self-reported time in a mood state (euthymic, depressed, or manic) would predict cerebellar volumes in the participants with bipolar disorder (**Supplemental Table 2**). No associations between these measures and cerebellar volumes were identified in this analysis (p > 0.05 and q > 0.05 for all cerebellar regions of interest).

### Effect of medication on cerebellar volume inferred from cross sectional assessment

Finally, we assessed whether taking a given class of psychotropic medication was associated with cerebellar volumes within bipolar disorder (**Supplemental Table 3**). For this analysis, we compared participants with bipolar disorder not taking a given class of medication to participants with bipolar disorder who were taking the class of medication. We did not analyze stimulant medication because use was infrequent (N = 13). We expected to observe differences in cerebellar volumes in those taking antipsychotics or lithium as compared to the other classes of medications based on previous reports, primarily in other brain areas (Fusar-Poli et al., 2013; Hallahan et al., 2011; Jones et al., 2022; Kim et al., 2013; Lyoo et al., 2010; Sun et al., 2018). Interestingly, the largest effects were instead observed with sedatives, a category that included benzodiazepines, sleep medications, barbiturates, and melatonin. Sedatives were associated with larger cerebellar volumes in left and right cerebellar white matter (left: p = 0.002, q = 0.008; right: p < 0.001, q = 0.003). Additionally, bipolar participants taking sedatives tended to have larger left and right cerebellar gray matter, but these were either not significant or did not survive correction (left: p = 0.054, q = 0.095; right: p = 0.024, q = 0.056). Demographic information on participants with bipolar disorder taking and not taking sedatives is available in **Supplemental Table 4**. To understand how these groups compared to control participants not taking sedatives, we evaluated the three groups using ANCOVAs, controlling for age and sex (**Figure 3; Supplemental Table 5**). In general, we found that control participants not taking sedatives and bipolar participants taking sedatives had similar non-vermal cerebellar volumes, while the bipolar participants not taking sedatives had smaller non-vermal volumes.

## Discussion

Here we report significantly smaller cerebellar cortex (gray matter) volumes bilaterally in individuals with bipolar disorder type I compared to controls. This difference did not extend to cerebellar white matter or vermal tissue and appeared diffuse throughout much of the cerebellar cortex. We found genetic loading for bipolar disorder did not explain the difference in gray matter, but that several variables were associated with cerebellar volumes within participants with bipolar disorder, including ACE score and current use of sedative medication. These findings add to the growing body of literature on cerebellar volumes in bipolar disorder and further highlight the need for nuanced study of the cerebellum as a potential player in bipolar disorder pathophysiology.

Our finding of reduced cerebellar hemisphere gray matter in bipolar disorder is consistent with several studies in the literature (de Zwarte et al., 2019; Lin et al., 2018; Moorhead et al., 2007; Sarıçiçek et al., 2015). It is important to note that our study sought to overcome potential limitations of some other studies in the literature by including larger sample sizes and specifying a single bipolar disorder subtype. Our bipolar disorder study’s sample size is second only to the multi-site ENIGMA consortium analysis, which also found smaller cerebellar gray matter volumes in bipolar disorder (de Zwarte et al., 2019). What might be causing the smaller cerebellar volumes remains unknown. One possibility is that the cerebellum may atrophy in bipolar disorder. A longitudinal study design would be needed to directly assess atrophy of the cerebellum, and our cross-sectional study was not designed to capture changes over time.

However, we did not observe correlation between duration of illness or symptom burden with cerebellar volumes, which would have been consistent with atrophy during disease course. It is possible that cerebellar volume changes occurred prior to diagnosis. This could indicate that bipolar disorder is associated with altered cerebellar development, which has been suggested to be especially sensitive to environmental factors due to protracted time course (Ciesielski and Knight, 1994). Consistent with this possibility, children and young adults at risk for bipolar disorder have different cognitive task activation patterns in the cerebellum (Johnsen et al., 2020) and are impaired on a cerebellar-dependent motor task (Giles et al., 2008). Furthermore, differences in white matter microstructure (Linke et al., 2020) and functional connectivity (Roberts et al., 2022) have been observed in the brains of youth at risk for bipolar disorder; however, the cerebellum has been understudied in this area. Further work is needed to examine the intersection of cerebellar development and bipolar disorder.

It is intriguing that bipolar PGS was not associated with changes in cerebellar gray matter volume while bipolar I diagnosis was. However, there is precedence for a psychiatric genome-wide polygenic risk scores to not correlate with brain volume differences, or for correlations to be very weak. For example, Reus and colleagues did not find any associations between polygenic risk score for bipolar disorder and any brain volumes examined; there were also no associations for major depression and only one association for schizophrenia (Reus et al., 2017). Similarly, other studies of schizophrenia found no associations between the significant mean differences in volume in schizophrenia and polygenic risk scores (Alnæs et al., 2019) and that genome-wide schizophrenia polygenic risk scores still had no associations with brain volumes even with increasing the sample size to almost 15,000 individuals (Grama et al., 2020). There are several possibilities for this type of discrepancy. One possibility is that the SNPs included in the calculation of the PGS do not capture the genetic differences responsible for reduced cerebellar gray matter. For example, if rare variants contribute to changes in cerebellar volume, these would not be included in the calculation of PGS, which could explain the lack of association. Another possibility is that the genetic underpinnings of reduced cerebellar gray matter are weaker predictors of bipolar disorder than other genes, and thus other genetic changes are predominantly determining the bipolar PGS and are thus drowning out the signal. It is possible that a more refined, non-genome-wide PGS would be better able to capture specific phenotypes, such as cerebellar volume changes; this has previously been suggested to be a better approach for finding brain structural changes in schizophrenia (Grama et al., 2020). They may also be environmental contributions resulting in structural changes, such as the reduced cerebellar gray matter that we observed in bipolar disorder. As environmental influences can change gene expression via epigenetic mechanisms, additional studies of alterations in gene expression in bipolar disorder would be beneficial. Finally, we cannot rule out complex interactions between certain genetic risk factors and other variables such as age and sex, or epistatic interactions between genetic variants; such complexities could mask true associations.

Given that bipolar PGS was not associated with cerebellar volumes across all participants, it is perhaps unsurprising that parental psychiatric history was also not associated with cerebellar volumes within bipolar disorder; however, it was intriguing that childhood adversity was associated with cerebellar white and gray matter volumes in bipolar disorder participants. This suggests that the ACE questionnaire may capture relevant information beyond familial mental illness, which could indicate an important role for environmental experiences in cerebellar development in bipolar disorder. There is precedence for ACEs to moderate other gray matter volume differences in bipolar disorder. In one study, participants with high ACEs showed reduced volumes in the inferior frontal gyrus bilaterally and the thalamus with bipolar disorder diagnosis, but not in participants with low ACEs (Poletti et al., 2016). This study showed reduced volume with increased ACEs, which is the opposite of what we observed in the cerebellum. However, the authors did not report on the cerebellum. A study outside of the context of bipolar disorder found smaller cerebellar gray matter volumes with increasing ACEs (Gehred et al., 2021). It is possible that the cerebellum changes differently from other regions with ACEs in bipolar disorder, potentially in a compensatory manner; however, further study in this area is required.

An important consideration in structural differences related to bipolar disorder involves medication exposure. In the present cross-sectional study, we found that sedative use in participants with bipolar disorder was significantly associated with larger cerebellar white matter volumes and non-significantly associated with larger cerebellar gray matter volumes. No other class of medications showed a significant relationship with cerebellar volumes. Thus, these findings would suggest that medications were not likely the cause of the reduced cerebellar volume when comparing participants with bipolar disorder to controls. However, we cannot completely rule out this possibility given the cross-sectional nature of the study. Prior studies have observed structural brain changes associated with antipsychotic and lithium treatment (Fusar-Poli et al., 2013; Hallahan et al., 2011; Jones et al., 2022; Kim et al., 2013; Lyoo et al., 2010; Sun et al., 2018). However, only one prior study (Huhtaniska et al., 2017) has been published related to the effects of sedatives on brain structure. This study found that higher cumulative benzodiazepine dose was associated with reduced caudate volume after controlling for cumulative antipsychotic dose and symptom severity. In the present study we observed an increase in cerebellar volume with participants being on a sedative. Furthermore, our study did not evaluate the effect of medications on the caudate since we were focused on factors that may influence our findings in the cerebellum. However, given these two sets of findings, one possible explanation is that greater cerebellar volumes result to compensate for caudate atrophy. An alternative possibility could be that our medication findings are a byproduct of confounding variables, such as differences in age. However, participants with bipolar disorder taking sedatives were older than those not taking sedatives, which would not explain why their cerebellar white matter volumes were larger. Yet another possibility could involve complex interactions between variables, such as those arising from polypharmacy, which this study did not address. Regardless of the underlying reasons, these findings raise the possibility that medication effects could diminish or mask true volume differences in bipolar disorder. Thus, cerebellar volume differences in bipolar disorder could be even greater without the confounds of medications. Future studies are needed to follow up on this possibility, and to investigate whether sedatives also associate with changes in cerebellar volumes in other psychiatric disorders.

The described work does come with some limitations. As mentioned, this study was cross-sectional, which limits our ability to understand how and when cerebellar differences emerge in bipolar disorder. This study design also limited analysis of medication effects, as we only examined current medications and not cumulative dosing. Furthermore, the presence of frequent polypharmacy was not addressed, and represents an important caveat to any observed medication effects. Finally, our sample included primarily White participants (88.7%), and thus our results may not generalize to all populations.

Overall, this paper supports the finding of smaller cerebellar cortices in bipolar disorder. Furthermore, it suggests that external sources such as medication exposure and childhood experiences may be important determinants of cerebellar volumes. Better understanding of cerebellar differences in bipolar disorder may help to advance understanding of bipolar disorder etiology, which in turn may help lead to novel preventative measures or therapeutic options.

