## Supplementary Material for "Cerebellar Morphological Differences in Bipolar Disorder Type I"

Supplemental Table 1. Many non-vermal cerebellar subregions from the SUIT atlas (Diedrichsen et al., 2009) were smaller in volume in bipolar disorder. Age and sex were included as covariates in all models. Significantly smaller ($q < 0.05$, *) and non-significantly smaller ($p < 0.05$ and $0.05 < q < 0.1$, ‡) areas are indicated.

| | Effect of bipolar diagnosis (N "Ctrl" = 81, N "BD" = 131) | | | |
|---|---|---|---|---|
| Cerebellar Volume (proportion of ICV) | Beta Coefficient Estimate | Estimated Mean Difference (converted to $cm^3$) | p-value | q-value |
| Lobules I-IV, left | -6.670e-05 | -0.098 | 0.1492 | 0.220 |
| Lobules I-IV, right | -4.909e-05 | -0.072 | 0.3329 | 0.388 |
| Lobule V, left | -1.599e-04 | -0.235 | 0.0045 | **0.032*** |
| Lobule V, right | -1.202e-04 | -0.177 | 0.0193 | **0.0498*** |
| Lobule VI, left | -2.006e-04 | -0.295 | 0.1038 | 0.162 |
| Lobule VI, right | -2.701e-04 | -0.397 | 0.0238 | 0.055‡ |
| Vermis VI | -5.722e-05 | -0.084 | 0.0305 | 0.061‡ |
| Crus I, left | -3.084e-04 | -0.454 | 0.0459 | 0.076‡ |
| Crus I, right | -1.990e-04 | -0.293 | 0.2443 | 0.311 |
| Vermis Crus I | 1.522e-07 | 0.000 | 0.3733 | 0.418 |
| Crus II, left | -3.070e-04 | -0.451 | 0.0130 | **0.0498*** |
| Crus II, right | -1.281e-04 | -0.188 | 0.2775 | 0.338 |
| Vermis Crus II | -2.928e-06 | -0.004 | 0.7567 | 0.785 |
| Lobule VIIb, left | -1.056e-04 | -0.155 | 0.0458 | 0.076‡ |
| Lobule VIIb, right | -1.226e-04 | -0.180 | 0.0282 | 0.061‡ |
| Vermis VIIb | 3.879e-07 | 0.001 | 0.9072 | 0.907 |
| Lobule VIIIa, left | -1.068e-04 | -0.157 | 0.0196 | **0.0498*** |
| Lobule VIIIa, right | -1.121e-04 | -0.165 | 0.0353 | 0.066‡ |
| Vermis VIIIa | -1.530e-05 | -0.023 | 0.4167 | 0.449 |
| Lobule VIIIb, left | -1.160e-04 | -0.171 | 0.0077 | **0.043*** |
| Lobule VIIIb, right | -1.160e-04 | -0.171 | 0.0151 | **0.0498*** |
| Vermis VIIIb | -1.131e-05 | -0.017 | 0.2004 | 0.267 |
| Lobule IX, left | -1.608e-04 | -0.236 | 0.0018 | **0.025*** |
| Lobule IX, right | -1.731e-04 | -0.255 | 0.0017 | **0.025*** |
| Vermis IX | -1.226e-05 | -0.018 | 0.1951 | 0.267 |
| Lobule X, left | -2.908e-05 | -0.043 | 0.0143 | **0.0498*** |
| Lobule X, right | -3.214e-05 | -0.047 | 0.0179 | **0.0498*** |
| Vermis X | -1.059e-05 | -0.016 | 0.0034 | **0.031*** |

Supplemental Figure 1. Representative brain with SUIT parcellation showing volume differences in bipolar disorder. Significantly smaller ($q < 0.05$, red) and non-significantly smaller ($p < 0.05$ and $0.05 < q < 0.1$, orange) areas are indicated.

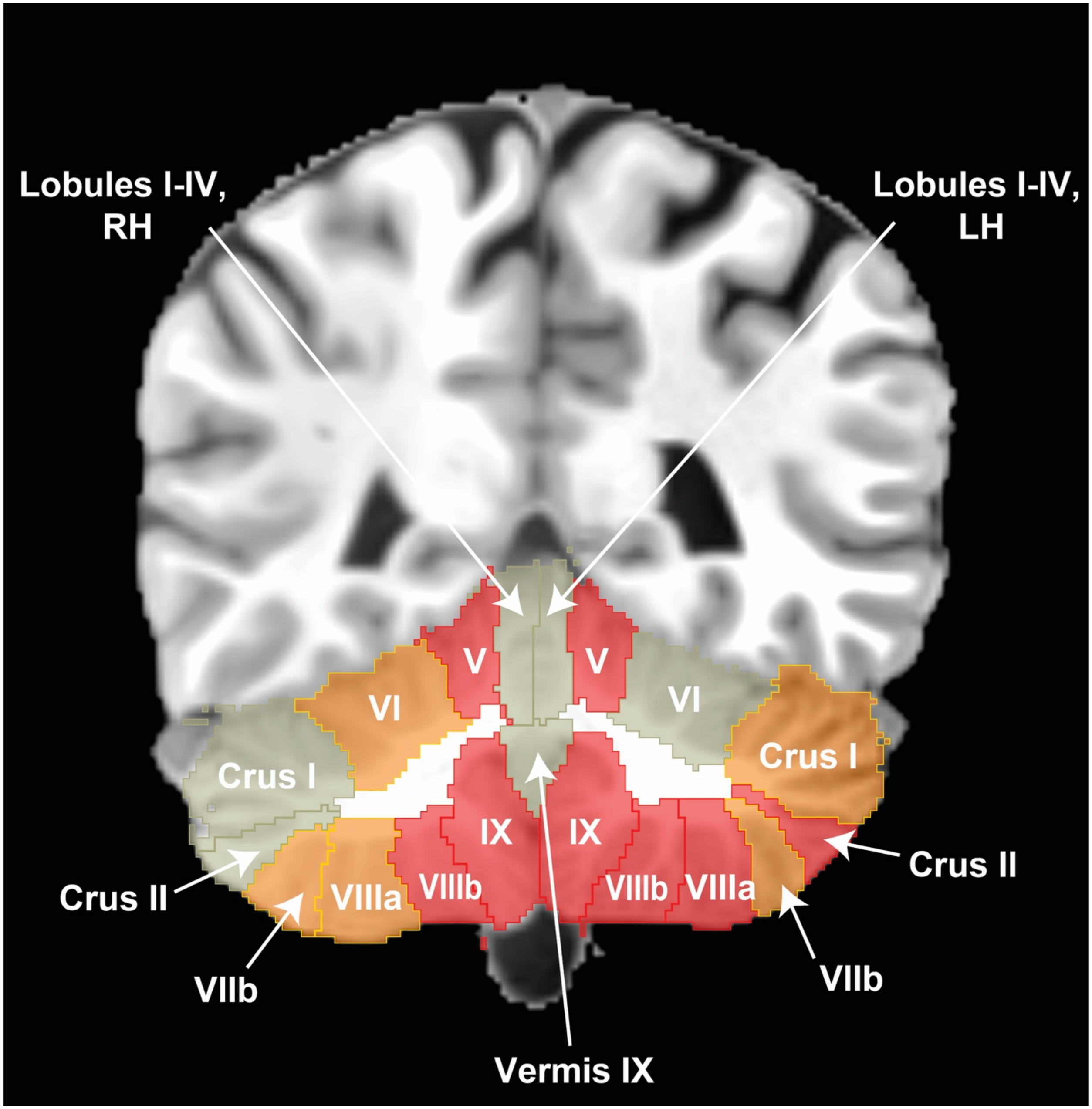

Supplemental Figure 2. Bipolar polygenic propensity scores (PGS) were higher in participants with bipolar disorder type I (t-test, $t(165) = -4.8564$, $p < 0.0001$). Control participants (Ctrl, N = 64) had a mean bipolar PGS of -0.431 (SD 0.979), while participants with bipolar disorder (BD, N = 103) had a mean bipolar PGS of 0.292 (SD 0.907).

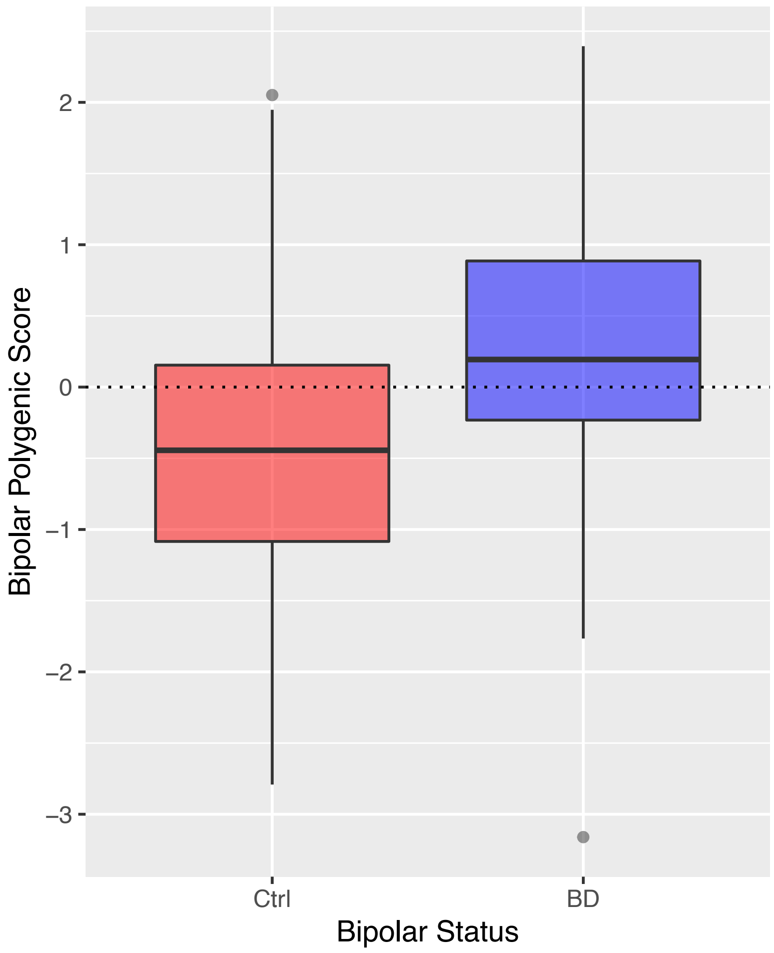

Supplemental Table 2. Illness duration and self-reported 10-year symptom burden were not associated with any cerebellar volumes in participants with BD. Age and sex were included as covariates in all models.

| Variable of interest (BD only) | Cerebellar Volume (prop. of ICV) | Beta Coefficient (SE) | p-value | q-value |
|---|---|---|---|---|
| Illness Duration (N = 131) | Left Cortex | -2.812e-05 (3.278e-05) | 0.393 | 0.877 |
| | Right Cortex | -3.037e-05 (3.270e-05) | 0.355 | 0.877 |
| | Left White Matter | 2.890e-06 (1.007e-05) | 0.775 | 0.877 |
| | Right White Matter | 7.495e-06 (9.937e-06) | 0.452 | 0.877 |
| | Vermis I-V | -7.748e-07 (4.983e-06) | 0.877 | 0.877 |
| | Vermis VI-VII | -9.729e-07 (2.115e-06) | 0.646 | 0.877 |
| | Vermis VIII-X | 5.383e-07 (2.892e-06) | 0.853 | 0.877 |
| Percent Time Euthymic (N = 129) | Left Cortex | 1.081e-05 (9.943e-06) | 0.279 | 0.548 |
| | Right Cortex | 8.543e-06 (9.938e-06) | 0.392 | 0.548 |
| | Left White Matter | 5.545e-07 (3.026e-06) | 0.855 | 0.915 |
| | Right White Matter | -3.209e-07 (2.986e-06) | 0.915 | 0.915 |
| | Vermis I-V | 1.427e-06 (1.504e-06) | 0.345 | 0.548 |
| | Vermis VI-VII | 9.719e-07 (6.386e-07) | 0.131 | 0.548 |
| | Vermis VIII-X | 7.666e-07 (8.660e-07) | 0.378 | 0.548 |
| Percent Time Depressed (N = 129) | Left Cortex | -6.223e-06 (1.145e-05) | 0.588 | 0.917 |
| | Right Cortex | -6.078e-06 (1.142e-05) | 0.596 | 0.917 |
| | Left White Matter | 6.109e-07 (3.472e-06) | 0.861 | 0.984 |
| | Right White Matter | 1.532e-06 (3.423e-06) | 0.655 | 0.917 |
| | Vermis I-V | -1.618e-06 (1.726e-06) | 0.350 | 0.917 |
| | Vermis VI-VII | -1.035e-06 (7.336e-07) | 0.161 | 0.917 |
| | Vermis VIII-X | -1.967e-08 (9.966e-07) | 0.984 | 0.984 |
| Percent Time Manic (N = 129) | Left Cortex | -1.677e-05 (1.652e-05) | 0.312 | 0.746 |
| | Right Cortex | -1.082e-05 (1.652e-05) | 0.514 | 0.746 |
| | Left White Matter | -2.808e-06 (5.019e-06) | 0.577 | 0.746 |
| | Right White Matter | -2.325e-06 (4.954e-06) | 0.640 | 0.746 |
| | Vermis I-V | -5.449e-07 (2.507e-06) | 0.828 | 0.828 |
| | Vermis VI-VII | -5.122e-07 (1.069e-06) | 0.633 | 0.746 |
| | Vermis VIII-X | -2.072e-06 (1.430e-06) | 0.150 | 0.746 |

Supplemental Table 3. Within participants with BD, sedative medication was significantly associated with larger cerebellar white matter bilaterally, with non-significantly larger cerebellar gray matter bilaterally. No other medication class survived FDR correction. Medication classes were assessed in separate models with age and sex as covariates. Significant results are indicated (q < 0.05, *).

| Variable of interest (BD only) | Cerebellar Volume (prop. of ICV) | Beta Coefficient (SE) | p-value | q-value |
|---|---|---|---|---|
| Antidepressants (N "No" = 55, N "Yes" = 76) | Left Cortex | -7.746e-04 (6.053e-04) | 0.203 | 0.362 |
| | Right Cortex | -6.471e-04 (6.053e-04) | 0.287 | 0.402 |
| | Left White Matter | -1.175e-05 (1.867e-04) | 0.950 | 0.950 |
| | Right White Matter | 4.643e-05 (1.845e-04) | 0.802 | 0.935 |
| | Vermis I-V | -1.466e-04 (9.143e-05) | 0.111 | 0.362 |
| | Vermis VI-VII | -4.945e-05 (3.897e-05) | 0.207 | 0.362 |
| | Vermis VIII-X | -1.015e-04 (5.284e-05) | 0.057 | 0.362 |
| Antipsychotics (N "No" = 58, N "Yes" = 73) | Left Cortex | 1.991e-05 (5.644e-04) | 0.972 | 0.972 |
| | Right Cortex | 2.112e-04 (5.630e-04) | 0.708 | 0.972 |
| | Left White Matter | 6.729e-06 (1.729e-04) | 0.969 | 0.972 |
| | Right White Matter | 3.003e-05 (1.709e-04) | 0.861 | 0.972 |
| | Vermis I-V | 1.008e-05 (8.555e-05) | 0.906 | 0.972 |
| | Vermis VI-VII | -2.463e-05 (3.627e-05) | 0.498 | 0.972 |
| | Vermis VIII-X | -5.205e-06 (4.965e-05) | 0.917 | 0.972 |
| Sedatives (N "No" = 75, N "Yes" = 56) | Left Cortex | 1.123e-03 (5.775e-04) | 0.054 | 0.095 |
| | Right Cortex | 1.310e-03 (5.733e-04) | 0.024 | 0.056 |
| | Left White Matter | 5.383e-04 (1.731e-04) | 0.002 | **0.008*** |
| | Right White Matter | 6.117e-04 (1.690e-04) | < 0.001 | **0.003*** |
| | Vermis I-V | 1.833e-05 (8.883e-05) | 0.837 | 0.837 |
| | Vermis VI-VII | 3.658e-05 (3.759e-05) | 0.332 | 0.388 |
| | Vermis VIII-X | 5.059e-05 (5.137e-05) | 0.327 | 0.388 |
| Anticonvulsants (N "No" = 75, N "Yes" = 56) | Left Cortex | 8.315e-04 (5.889e-04) | 0.160 | 0.321 |
| | Right Cortex | 7.470e-04 (5.887e-04) | 0.207 | 0.321 |
| | Left White Matter | 2.185e-04 (1.808e-04) | 0.229 | 0.321 |
| | Right White Matter | 2.236e-04 (1.787e-04) | 0.213 | 0.321 |
| | Vermis I-V | 4.506e-05 (8.989e-05) | 0.617 | 0.617 |
| | Vermis VI-VII | 8.518e-05 (3.746e-05) | 0.025 | 0.172 |
| | Vermis VIII-X | 5.042e-05 (5.203e-05) | 0.334 | 0.390 |
| Lithium (N "No" = 95, N "Yes" = 36) | Left Cortex | -5.768e-04 (6.249e-04) | 0.358 | 0.626 |
| | Right Cortex | -4.119e-04 (6.248e-04) | 0.511 | 0.715 |
| | Left White Matter | -2.712e-04 (1.906e-04) | 0.157 | 0.551 |
| | Right White Matter | -2.050e-04 (1.891e-04) | 0.280 | 0.626 |
| | Vermis I-V | -4.718e-05 (9.496e-05) | 0.620 | 0.724 |
| | Vermis VI-VII | -4.234e-06 (4.037e-05) | 0.917 | 0.917 |
| | Vermis VIII-X | 1.091e-04 (5.431e-05) | 0.047 | 0.327 |

Supplemental Table 4. Demographic breakdown of participants with BD depending on sedative medication status. Significant differences are indicated ($p < 0.05$, *).

| Variable | | Not taking sedatives (N=75) | Taking sedatives (N=56) | p value |
|---|---|---|---|---|
| Biological Sex | (F, M) | (45, 30) | (40, 16) | 0.175 |
| Age (years) | Mean (SD) | 37.068 (13.253) | 43.457 (12.937) | 0.007* |
| BMI | Mean (SD) | 29.097 (6.905) | 33.434 (8.238) | 0.001* |
| Educational Attainment | Mean (SD) | 14.827 (2.165) | 14.964 (2.182) | 0.720 |
| Self-report SES | Mean (SD) | 5.453 (1.742) | 4.927 (2.159) | 0.127 |
| | N missing | 0 | 1 | |
| Intracranial Volume (cc) | Mean (SD) | 1499.461 (192.659) | 1447.337 (168.087) | 0.108 |
| Any parental psych illness (Yes/No) | (Y, N) | (51, 17) | (41, 9) | 0.365 |
| | N "Unknown" or missing | 7 | 6 | |
| ACE Score | Mean (SD) | 3.464 (2.697) | 4.075 (2.654) | 0.274 |
| | N missing | 19 | 16 | |
| MADRS Score | Mean (SD) | 12.947 (9.744) | 16.500 (8.534) | 0.031* |
| | Range | 0 - 32 | 1 - 44 | |
| YMRS Score | Mean (SD) | 5.173 (5.168) | 8.714 (8.405) | 0.003* |
| | Range | 0 - 22 | 0 - 40 | |
| BAI Score | Mean (SD) | 10.838 (10.539) | 16.870 (12.038) | 0.003* |
| | N missing | 1 | 2 | |
| | Range | 0 - 40 | 1 - 45 | |
| Medication Status | | | | |
| Antidepressant(s) | N Yes (% Yes) | 34 (45.3%) | 42 (75.0%) | < 0.001* |
| Antipsychotic(s) | N Yes (% Yes) | 40 (53.3%) | 33 (58.9%) | 0.524 |
| Anticonvulsant(s) | N Yes (% Yes) | 23 (30.7%) | 33 (58.9%) | 0.001* |
| Stimulant(s) | N Yes (% Yes) | 5 (6.7%) | 8 (14.3%) | 0.149 |
| Lithium | N Yes (% Yes) | 20 (26.7%) | 16 (28.6%) | 0.809 |
| Time from first mood (BD) | Mean(SD) | 16.253 (13.003) | 21.196 (13.127) | 0.034* |
| | Range | 0.000 - 43.000 | 0.000 - 46.000 | |
| % time euthymic (BD) | Mean (SD) | 47.9 (28.4) | 34.4 (27.8) | 0.008* |
| | Range | 0 - 98% | 0 - 98% | |
| | N missing | 0 | 2 | |
| % time depressed (BD) | Mean (SD) | 34.1 (23.8) | 45.5 (26.1) | 0.011* |
| | Range | 0 | 2 | |
| | N missing | 0 - 80% | 2 - 90% | |
| % time manic (BD) | Mean (SD) | 18.0 (16.1) | 20.1 (18.8) | 0.486 |
| | Range | 1 - 93% | 0 - 95% | |
| | N missing | 0 | 2 | |
| Bipolar PGS | Mean (SD) | 0.410 (1.006) | 0.126 (0.726) | 0.117 |

Supplemental Table 5. ANCOVA comparisons of cerebellar volumes in control participants not taking a sedative (Ctrl (-)), participants with BD not taking a sedative (BD (-)), and participants with BD taking a sedative (BD (+)). Age and sex were used as covariates. There were significant effects of group bilaterally for both cerebellar gray matter and white matter ($q < 0.05$, *). Pairwise comparisons with Tukey correction suggested the BD (-) group tended to have smaller volumes than the other two groups (Tukey-adjusted $p < 0.05$, *).

| Variable of interest | Cerebellar Volume (prop. of ICV) | ANCOVA p-value | ANCOVA q-value | EMM Group comparisons (Tukey-adj. p-values) | | |
|---|---|---|---|---|---|---|
| | | | | Ctrl (-) vs BD (-) | Ctrl (-) vs BD (+) | BD (-) vs BD (+) |
| Sedative group (N "Ctrl (-)" = 76, N "BD (-)" = 75, N "BD (+)" = 56) | Left Cortex | 0.001 | 0.004* | **0.001*** | 0.515 | 0.071 |
| | Right Cortex | 0.002 | 0.004* | **0.002*** | 0.796 | **0.041*** |
| | Left White Matter | 0.002 | 0.004* | **0.018*** | 0.733 | **0.004*** |
| | Right White Matter | 0.001 | 0.004* | **0.034*** | 0.352 | **0.001*** |
| | Vermis I-V | 0.859 | 0.859 | 0.851 | 0.988 | 0.936 |
| | Vermis VI-VII | 0.157 | 0.183 | 0.139 | 0.833 | 0.475 |
| | Vermis VIII-X | 0.130 | 0.183 | 0.119 | 0.874 | 0.390 |
